# Microcircuitry of Performance Monitoring

**DOI:** 10.1101/187989

**Authors:** Amirsaman Sajad, David C. Godlove, Jeffrey D. Schall

## Abstract

**Acknowledgments:** This work was supported by R01-MH55806, P30-EY08126, and by Robin and Richard Patton through the E. Bronson Ingram Chair in Neuroscience. We thank J. Easley M. Feurtado, M. Maddox, P. Middlebrooks, S. Motorny, J. Parker, M. Schall, C.R. Subraveti, A. Tomarken, and L. Toy for animal care and other technical assistance. We thank J. Brown, M. Cox, K. Dougherty, S. Errington, A. Maier, V. Stuphorn, A. Tomarken, and J. Westerberg for helpful discussions and comments on the manuscript.

**Conflicts of interest:** None

Cortical circuit mechanisms in medial frontal cortex enabling executive control are unknown. Hence, in monkeys performing a saccade countermanding task to earn larger or smaller fluid rewards, we sampled spiking and synaptic activity simultaneously across all layers of the supplementary eye field (SEF), an agranular cortical area contributing to performance monitoring in nonhuman primate and human studies. Laminar-specific synaptic currents with associated spike rate facilitation and suppression represented error production, reward gain or loss feedback, and reward delivery. The latency, polarity and magnitude of current and spike rate modulation were not predicted by the canonical cortical microcircuit. Pronounced synaptic currents in layer 2/3, which are modulated by loss magnitude, will contribute to the error-related negativity (ERN) and feedback-related negativity (FRN). These unprecedented findings reveal critical features of the cortical microcircuitry supporting performance monitoring and demonstrate that SEF can contribute to the error‐ and feedback-related negativity.

**Subject terms:** countermanding, stop signal task, goal selection, response inhibition, executive control, canonical cortical microcircuit, error-related negativity, reinforcement learning, reward prediction error

## Introduction

That the medial frontal lobe contributes to performance monitoring and executive control is beyond dispute, but the specific mechanisms remain uncertain because diverse findings have supported divergent, even incompatible hypotheses (Heilbronner & Hayden 2016; Kolling et al. 2016; Shenhav et al. 2016; Stuphorn 2015; Procyk et al. 2016). Noninvasive measures used to investigate this question are the error-related negativity (ERN) and feedback-related negativity (FRN), event-related potentials indexing performance monitoring in humans (reviewed by Gehring et al. 2012), which have been found in macaque monkeys (Vezoli & Procyk 2009; Godlove et al. 2011; Phillips & Everling 2014). Intracortical recording of spiking and LFP signals are necessary to resolve the nature and sequence of synaptic currents and spike rate modulation across cortical layers, which will contribute to resolution of major alternative hypotheses.

Performance monitoring and executive control is investigated with the stop-signal (countermanding) task (Schall & Boucher, 2007; Verbruggen & Logan 2009). Macaque monkeys performing saccade countermanding strategically adapt saccade latency according to the trial history (Emeric et al. 2007; Nelson et al. 2010). Multiple studies in different laboratories have demonstrated contributions of the supplementary eye field (SEF), an agranular area on the dorsomedial convexity in macaques, to performance monitoring and executive control. SEF neurons signal errors and reinforcement in spikes (Stuphorn et al. 2000) and local field potentials (Emeric et al. 2010). Intracranial recordings in humans show performance monitoring signals in the adjacent supplementary motor area (Bonini et al. 2014). Similar signals are found in dorsal ACC (Ito et al. 2003; Nakamura et al. 2005; Amiez et al. 2006; Seo & Lee 2007; Emeric et al. 2008; Kennerley et al. 2011; Ebitz & Platt 2015; Michelet et al. 2015). SEF also signals proactive inhibition predicting whether movements will ultimately be inhibited (Stuphorn et al. 2010). Finally, subthreshold electrical stimulation of SEF improves countermanding performance by increasing the duration of the GO process (Stuphorn & Schall 2006), accomplished by delaying the onset of accumulation of pre-movement activity in FEF and SC (Pouget et al. 2011).

Much evidence shows that computations to transform representations occur through a canonical cortical microcircuit (CCM) (Bastos et al. 2012). Although formulated based on information from sensory cortical areas, the circuit in agranular areas like SEF is different (Shipp 2005; Weiler et al. 2008; García-Cabezas and Barbas, 2014; Godlove et al. 2014; Beul and Hilgetag 2015; Ninomiya et al. 2015) but evidently suited for performance monitoring (Cohen 2014). However, nothing is known about the laminar distribution of synaptic currents and spike rate modulation in a medial frontal area of monkeys during a cognitively demanding task. Therefore, we obtained samples of spikes and synaptic potentials across all layers of SEF during performance of saccade countermanding. The data provide the first detailed outline of the microcircuitry of SEF supporting performance monitoring. The findings show how error as well as reward gain and loss signals arise within and flow across layers. The results will contribute to resolution of alternative hypotheses about medial frontal function and show how SEF can contribute to the ERN and FRN.

## Results

### Countermanding performance with asymmetric reinforcement

Neural data was recorded from two macaque monkeys performing the saccade countermanding task (Hanes & Schall 1995) with asymmetric fluid reward volumes and explicit success feedback tone cues (Fig. 1). Both monkeys exhibited sensitivity to the stop signal and to reward magnitude (Supplementary Fig. 1, Table S1). The probability of failing to cancel the saccade on stop trials increased with stop signal delay. Response time (RT) was significantly shorter in non-canceled compared to no stop signal trials, and on high compared to low reward magnitude trials. Stop signal reaction time (SSRT) was of typical magnitude for this task and was unaffected by reward magnitude. RT was delayed on no stop signal trials following both canceled and non-canceled stop signal trials relative to no stop trials.

**Figure 1.**
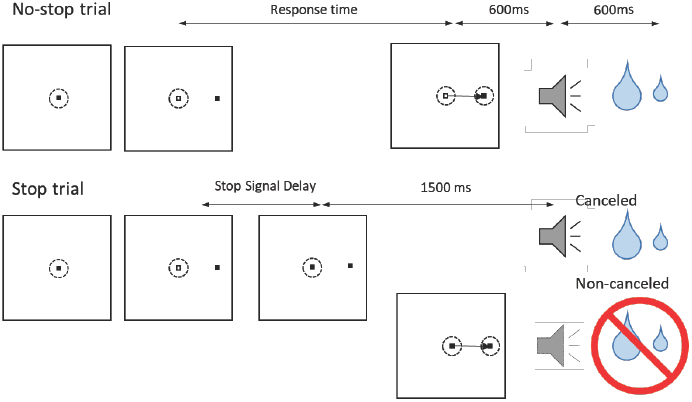
Asymmetrically rewarded saccadic stop-signal task. Trials were initiated when monkeys fixated a central point. After a variable time, the center of the fixation point was extinguished leaving an outline. A peripheral target was presented simultaneously at one of two possible locations. The location of the target cued the monkey that either a large or small magnitude reward could be obtained on the current trial. These reward-to-location mappings reversed predictably in blocks of 10-30 correct no-stop trials. On No-stop trials monkeys were required to make a saccadic eye movement towards the target. 600 ms following the execution of the correct saccade the monkey was presented with a high-pitch auditory feedback tone, and 600 ms later fluid reward was provided. On stop signal trials, after the target appeared, the center of the fixation point was re-illuminated after a variable time (stop signal delay) instructing the monkey to cancel the saccade. In ∼50% of the trials the monkey successfully stopped the saccade (canceled), the same high-pitch auditory tone was presented after a time delay and juice reward was provided. In the other ∼50% of trials the monkey made an error saccade (noncanceled), and a low-pitch feedback tone was presented 600 ms after the saccade, and no reward was delivered. Note that the correct (no-stop) and error (non-canceled) saccade trials showed a similar temporal sequence of events.

### Location of samples

SEF is located in the dorsal medial convexity in macaques, making it readily accessible for laminar electrode array recordings perpendicular to the cortical layers. SEF was located with anatomical landmarks and intracortical electrical microstimulation. Using linear microelectrode arrays (Plexon U-probe, 150 μm inter-contact spacing), we recorded spikes and LFPs from SEF of 2 macaque monkeys. We acquired 33816 trials (Eu: 11583, X: 22233) in 29 sessions (Eu: 12, X: 17). Across all recording sessions we isolated 575 single units (Eu: 331, X: 244) of which 61 (Eu: 51, X: 10) were modulated when countermanding errors were produced and 265 (Eu: 105, X: 160) were modulated when reinforcement gain or loss was cued or delivered. Of these, 69 neurons multiplexed signal types (Supplementary Fig. 2a; Table S2). In 16 sessions, recordings obtained with electrode arrays oriented perpendicular to the cortical layers, verified through combined MR and CT imaging (Fig. 2; Godlove et al. 2014), sampled 293 neurons of which 173 (Eu: 65/104 neurons; X: 108/189) contributed to these results (Table S2).

Sampling depths were aligned across sessions (Supplementary Fig. 2b). Neural signals were sampled across multiple sessions from the same coordinates to investigate reproducibility (Supplementary Fig. 2c). Signals were assigned to SEF layers that were assessed in a battery of histological sections visualized with Nissl (Matelli et al. 1991), neuronal nuclear antigen, Gallyas myelin, cytochrome oxidase, acetylcholinesterase, nonphosphorylated neurofilament H (Geyer et al. 2000), parvalbumin, calbindin, and calretinin. Further information about laminar structure was assessed through the pattern of cross-frequency phase-amplitude coupling across the depth of SEF (Ninomiya et al. 2015). Due to noise in the estimates, some units were assigned to L1 or deeper L6.

**Figure 2.**
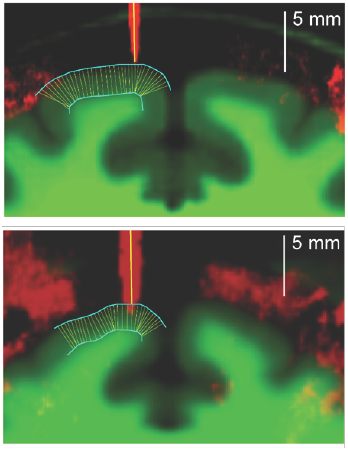
Recording track of two perpendicular penetrations. Coronal planes of the SEF and surrounding gray matter (dark green) and white matter (light green) tissue is shown for monkey Eu (top) and monkey X (bottom). Thick vertical yellow line shows trajectory of linear electrode array in guide tubes for two of the three perpendicular penetrations. The red vertical line shows the guide tube orientation for two perpendicular penetrations. Cyan lines show pial surface and transition from gray matter to white matter. Thin yellow lines show the result of an automated algorithm that minimized distance between the pial surface and gray matter to calculate angles perpendicular to gray matter. At these penetration sites the thick and think yellow lines almost perfectly align. Modified from Godlove et al. (2014).

### Error signals

#### Spiking activity

By design, monkeys produce non-canceled errors on ∼50% of stop signal trials. Error-related spiking activity was identified by comparing discharge rates between errant non-canceled and correct no stop trials (Fig. 3a). Error-related spiking was observed in multiple penetrations in both monkeys, but such activity was noticeably concentrated in particular locations (Chi square contingency test of incidence across penetration locations, *χ*^2^(9, *N* = *575*) = 101.525, p << 0.001; Supplementary Table 2). The difference in prevalence of error-related activity was not dependent on monkey identity or performance.

**Figure 3.**
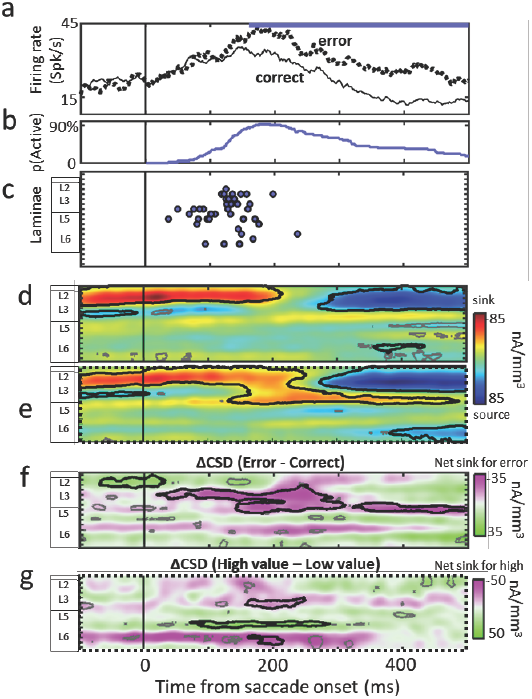
Temporal and Laminar organization of error-related modulations in SEF. **a)** Example error-related spiking activity aligned on saccade onset of a single neuron with elevated discharge rate on error (non-canceled) relative to correct (no-stop) trials. Blue bar represents the duration of significant error activity in this unit. **b)** Proportion of units signaling error as a function of time. A monotonic recruitment of error-units after error saccade was followed by decreasing active neurons. **c)** Latency of error-related modulation as a function of cortical depth in channel units (150 μm). Earliest error-related activity is observed in L5 followed by modulation in L3 and L6. **d)** Grand average current source density (CSD) for correct (top) and error (bottom) conditions. Black contours represent patches of significant current sink or source. While the CSD patterns show a similar extended current sink (red) in superficial layers, a later sink in L3 and L5 is observed on error trials. This difference is highlighted in the grand average ΔCSD (error – correct) as a net sink in current in L3 and L5 (in purple) associated with errors **(e)**. **f)** Difference CSD (ΔCSD) obtained from CSD on error trials associated with high cost vs. low cost, showing differential synaptic current flow for different error cost in L3 upper and lower L6.

The characteristics of the neurons corresponded to previous descriptions (e.g., Stuphorn et al. 2000). Most of the error-related neurons (45/61 neurons) were not lateralized, and neurons with differential contra‐ and ipsiversive activity showed similar patterns of activity. Error-related modulation across the sampled cells on average ± SD persisted for 253 ± 155 ms. The proportion of recruited error-related neurons monotonically increased from 40 ms, peaked at ∼190 ms then declined to about 30% after 400 ms (Fig. 3b). There was no significant difference in duration between modulations in different layers and early‐ and late-onset modulations. Error-related spiking did not predict post-error slowing.

We defined a *reward sensitivity index* to characterize whether spike rates were modulated by reward magnitude (RSI = (Reward_High_ – Reward_Low_)/ (Reward_High_ + Reward_Low_)). Notably, most error neurons exhibited larger responses when high relative to low reward was lost (i.e., RSI > 0; 43/61 neurons, paired Wilcoxon test across sample, df = 60, p = 3.4 x 10^−5^, Supplementary Fig. 3a). These neurons were invariably distributed across all cortical laminae, and we found no clear relationship between the timing and RSI of activity.

The laminar profile of error-related spiking could be determined for 42 of 61 neurons. Error-related spiking appeared earliest after saccade initiation in deep L3 and L5 in the 100 ms after the error (Fig. 3c). Error-related spiking then arose in upper L2/3 and L6. Spike widths (trough to peak duration) of error neurons in L2/3 were significantly narrower than those in L5/6 (unpaired two-sample Wilcoxon test, df = 41, p = 0.022, Supplementary Fig. 3b).

#### Field potentials

The polarization and time course of LFP replicate previous findings (Emeric et al. 2010). On both correct no-stop and error non-canceled trials a negative polarization was associated with the saccades. Errors were associated with greater negative polarization. The error-related LFP negativity occurs at all cortical depths, arising earliest in L3, L5 and upper L6 and progressively later in more superficial and deeper layers. The negativity is briefest (<100 ms) in superficial layers and longest (>200 ms) in L3 and below (Supplementary Fig. 3c). The time course of the net negativity in grand average LFPs paralleled the onset of error-related spiking activity and the recruitment of error neurons. Like the ERN, LFP polarization magnitude was modulated by the value of error cost (Supplementary Fig. 3d). Larger error cost was associated with a significant, brief (∼100 ms) net negativity most prominent in L2/3 and L5 followed by a brief net positivity (∼100 ms) most prominent in L1, L2, L5 and L6.

#### Laminar current density

CSD was extracted from LFPs to determine the spatiotemporal flow of current in the SEF. Previously, we showed that light flashes evoke laminar specific CSD in SEF (Godlove et al. 2014). We now describe task-related CSD in SEF with duration far exceeding the brief post-stimulus sensory epoch considered by most studies (e.g., Maier et al., 2011). CSD on correct trials aligned to saccade initiation exhibited a current sink in L2/3 beginning before the saccade and continuing ∼200ms thereafter (Fig. 3d). This is unrelated to saccade generation *per se* because no current flow is observed associated with spontaneous saccades in the dark (Godlove et al. 2014). On error trials, the peri-saccadic sink in L2/3 occurs which is followed by a significant current sink in deeper L3 and upper L5 (Fig. 3e).

To characterize the error-related CSD, we subtracted the CSD on correct from that on error trials (Fig. 3f). The grand average ΔCSD shows that the errors are associated with stronger current sinks initially in L3 ∼30-130 ms after the error, then in L3 and L5 from ∼150-300 ms, and finally only in L5. Comparison of CSD pattern between high-cost vs. low-cost error trials showed a net negativity in current in layers L3 and L6 and a net positivity in current in layers L6 associated with larger cost (Fig. 3g). These differential CSD patterns were observed ∼100-250 ms after the saccades, coincident with significant error-related current sinks in L3 and L5, and the negativity in field polarizations.

### Reinforcement signals

Each saccade was followed, after 600 ms, by an auditory feedback tone distinguishing correct and errors trials. On correct trials juice reward was delivered 600 ms after the tone. This temporal structure dissociated self-generated signals (like expectation of fluid reward) from responses to sensory cues (like consumption of fluid). This section describes the functional architecture of reinforcement-related processing in SEF.

#### Spiking activity

Reinforcement-related spiking activity was identified by comparing discharge rate between unrewarded and rewarded trials (Fig. 4a,b). The measurement began at the presentation of the feedback tone until 200 ms following scheduled delivery of the reward. Any neuron with significant modulation starting in this period was considered reinforcement-related. Neurons signaling feedback and reward delivery were observed in both monkeys and all recording sites (Table S2). The depth distribution of reinforcement-related neurons was surprisingly consistent across sessions and recording sites in both monkeys (Supplementary Fig. 2d).

**Figure 4.**
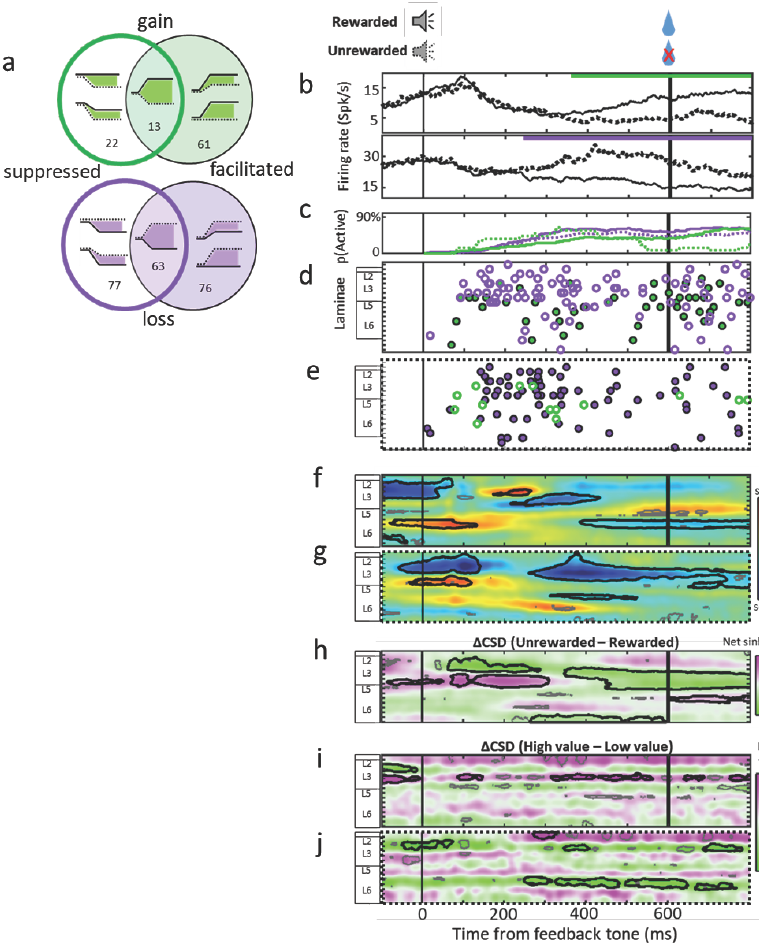
Temporal and laminar organization of reinforcement-related modulations in SEF. **a)** Venn diagram depicting counts of different reinforcement-related modulation types. Inside the circles schematics of possible modulations on rewarded (solid line) and unrewarded (dotted line) trials are shown. **b)** Example reinforcement-related spiking activity aligned on feedback tone of two units signaling reward gain (top) or reward loss (bottom). Green and purple lines represent the period of significant differential activity for the units. **c)** shows the time course of the proportion of active units exhibiting facilitation (solid line) or suppression (dashed line) signaling gain (green) or loss (purple). **d-e)** The latency of reinforcement-related activity as a function of cortical depth is plotted for rewarded (d) and unrewarded trials **(e)**. Facilitated gain units (filled green circle) and suppressed loss units (open purple circles) respond to positive outcomes while facilitated loss units (filled purple circle) and suppressed gain units (open green circle) respond to negative outcomes. **f-g)** Grand average CSD for rewarded **(f)** and unrewarded **(g)** trials with similar conventions as in Fig3d-e. **h)** ΔCSD obtained from difference between rewarded and unrewarded CSDs. **i-j)** ΔCSD obtained from difference in CSD on high‐ and low-value rewarded **(i)** and unrewarded **(j)** trials (purple: net current sink for high value). The sinks in current associated with gain or loss do not show dynamic interlaminar flow through time.

The characteristics of the reinforcement-related modulation corresponded to previous descriptions (e.g., Stuphorn et al. 2000; So and Stuphorn, 2010, Seo and Lee, 2009). Neurons were distinguished based on the time, sign, and valence of modulation (Fig. 4a). Gain neurons (88 neurons with 96 modulation intervals) exhibited higher discharge rate on rewarded than on unrewarded trials. This could result from either facilitation on rewarded trials (61 intervals), suppression on unrewarded trials (22 intervals), or both (13 intervals). Loss neurons (184 neurons with 216 modulation intervals) exhibited higher discharge rate on unrewarded than on rewarded trials. This could result from either facilitation on unrewarded trials (76 intervals), suppression on rewarded trials (77 intervals), or both (63 intervals). On correct trials, facilitated gain (68 neurons, 74 intervals) was less common than suppressed loss (117 neurons, 140 intervals). On unrewarded error trials, facilitated loss (129 neurons, 139 intervals) was more common than suppressed gain (34 neurons, 35 intervals). Thus, SEF spiking increases more for negative than positive outcomes. Further, the modulation on rewarded trials was sensitive to reward amount (RSI of suppressed loss responses < 0, One-sample Wilcoxon test, df = 139, p < 0.0073), especially in neurons with modulation onset <250 ms after the tone (Supplementary Fig. 4a).

The onset time and duration of reinforcement-related spiking modulation was quite variable. Notably the onset of the activity was not briskly synchronized on tone or reward, which suggests that it arises from intrinsic processing in the SEF rather delivered as a response to the sensory aspects of these events. Overall, the proportion of active gain and loss neurons monotonically increased until 400 ms after feedback delivery. This proportion was sustained for loss neurons until after the fluid reward was delivered and only moderately increased for excited gain neurons after juice delivery. In contrast, the proportion of recruited suppressed gain neurons dropped ∼200 ms before reward delivery (Fig. 4c).

In samples from penetrations perpendicular to the layers we found reinforcement-related neurons in all layers (Fig 4d,e), but gain and loss neurons showed significantly different depth distributions (Table S2). Loss neurons were least common in L5. Suppressed loss neurons with earliest modulation onset were most common in L2/3 but after ∼250 ms they were also observed in deep L6. Facilitated loss neurons were also common in upper L2/3; most of these were also suppressed on rewarded trials. Facilitated loss neurons with no modulation on rewarded trials however were more common in L6. Gain neurons were common in deep L3, L5, and L6 with equal distributions of facilitated and suppressed examples.

Spike width in SEF varies with layer and putative neuron type (Godlove et al. 2014). Spike width of both facilitated and suppressed L2/3 neurons were significantly narrower than those in L5/6 (F(1,307) = 12.9, p < 0.001) (Supplementary Fig. 4b). Loss neurons in L2/3 and L5/6 had spikes <250 μs wide, consistent with arising from parvalbumin interneurons.

#### Field potentials

Feedback and reinforcement-related field potentials were analyzed in the interval between the tone and 200 ms after scheduled delivery of reward. LFPs exhibited an initial negativity on most channels in response to the tone on both rewarded and unrewarded trials (Emeric et al., 2010). This sensory-related polarization was followed by a positivity that was more pronounced on unrewarded trials (Supplementary Fig. 4c). Relative to unrewarded trials, the LFP on rewarded trials showed a significant net negativity at all depths that started before the feedback tone and lasted until ∼100ms after the tone. The polarization difference then shifted to a net negativity on rewarded trials that reached significance ∼250 ms before juice delivery and lasted until the end of the analyzed period (200ms after juice delivery).

LFP polarization was also modulated by the magnitude of reward gain or loss. Relative to trials with larger reward, the LFP on trials with smaller reward exhibited greater negativity in the interval around the feedback tone followed by a significant net negativity for larger reward in the 200 ms interval preceding juice delivery. This negativity had different temporal structure at different depths, with shortest duration in L2, but it was consistently observed within the 200 ms pre-juice period and ended before or at juice delivery time (Supplementary Fig. 4d). The value of reward loss was signaled by a brief, significant net negativity associated with larger loss that appeared ∼200 ms after the auditory feedback cue, in lower layers corresponding to L3, L5, and L6 (Supplementary Fig. 4e).

#### Laminar current density

The grand average CSD on rewarded no-stop trials and unrewarded non-canceled trials exhibited different spatiotemporal profiles (Fig. 4). On rewarded trials, a current sink was observed in L6 beginning <100ms before the positive feedback tone, reaching its peak magnitude ∼80ms after the tone and persisting until 150ms after the tone. This was followed by a current sink in L3 ∼200-300ms after the auditory feedback. On unrewarded trials, while there was an overall negativity in current in L6 in the early period following the tone, it did not reach significance; instead, a significant sink in current was observed in L3, similar in time course to the early sink observed in L6 on rewarded trials. Unlike rewarded trials, on unrewarded trials no significant sinks in current were observed beyond 100 ms after the tone.

The reinforcement-related ΔCSD was determined by subtracting the CSD on rewarded trials from that on unrewarded trials (Fig. 4h). Reward loss was associated with significantly greater current negativity in L3, from ∼80 to 300 ms after the tone. On the other hand, reward gain was associated with greater current negativity during the same interval more superficially in L2/3, though the early phase of this difference period overlapped with a large current source on unrewarded trials, which limits its interpretations. Also, reward gain was associated with greater current negativity from ∼250 ms before juice delivery to the 200 ms post-reward limit of analysis. This net negativity in current flow on rewarded trials was coincident with the significant negativity in field potentials.

Next we compared the current flow for high vs. low magnitude of gain and loss. Higher reward gain was associated with a greater current negativity in the L3 current sink associated with reward gain. This negativity persisted intermittently from ∼100ms after the auditory tone until 200 ms after juice delivery, most prominently during the 250ms before reward delivery. Although the weak sink in current in L6 on unrewarded trials did not reach significance, it showed sensitivity to reward amount. Less reward loss was associated with greater current negativity in the L6 sink. This negativity also persisted intermittently from 300 ms before to 200 ms after scheduled juice delivery. A brief net negativity in L3 was also observed, though this difference arose from differences in current source magnitudes, which are less certain in interpretation. Surprisingly and importantly, the modulations in CSD related to magnitude of gain/loss involved no inter-laminar current flow.

### Laminar distribution of of error and reinforcement spiking

Fig. 5 summarizes the laminar distribution of neurons signaling error, gain and loss through facilitation and suppression in all of the penetrations perpendicular to the layers. The incidence of neurons through depth was scaled by the overall distribution of sampled neurons. Neurons with different patterns of modulation were not distributed uniformly across the layers. Error-responsive neurons were most common in lower L3, upper L5, and lower L6. Neurons signaling loss of reward were most common across the sample, and they were most concentrated in L2 and L6, less common in L3 and nearly absent in L5. Neurons facilitated or suppressed by loss of reward had similar laminar distributions. Neurons signaling gain were less common overall, and those recorded were most common in lower L3 and L5. Gain neurons were more often facilitated than suppressed by reward cue or delivery. Thus, the following trends were noted: In L2 and upper L3 most neurons signal reward loss, both via suppression to positive events and facilitation to negative events. In lower L3 neurons signal error, loss and gain. Neurons in L5 were more likely to signal error and gain. In L6 most neurons signaled error or loss, most likely via facilitation.

**Figure 5.**
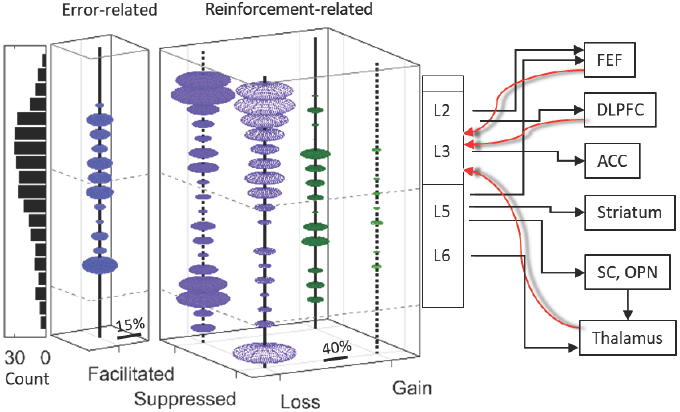
Functional architecture of SEF for performance monitoring. Left, distribution of units sampled across depth, pooled across all perpendicular penetrations. Center, proportion of error-related and reinforcement-related neurons normalized by sample density. Error-related neurons were densest in lower L3,L5 and lower L6. Gain neurons were found mainly in lower L3 and lower L5 and L6. Loss neurons were most frequent in upper L2/3, infrequent in L5, and more frequent in L6. Facilitation was more frequent than suppression in L6. Some loss neurons were putative parvalbumin neurons, but all others had wide spikes characteristic of pyramidal projection neurons. Thus, the proportions of various signals arise through intrinsic processing of laminar-specific afferents and influence different structures through laminar-specific efferents. Right, selected outputs (black) and inputs (red) across SEF layers. Further details in text.

## Discussion

We provide the first description of the laminar distribution of error and reward signals in the medial frontal cortex of primates. Patterns of unit discharges and LFP polarization signaling error, loss and gain replicated previous studies of SEF during saccade countermanding (Stuphorn et al. 2000; Emeric et al. 2010) and other tasks (So & Stuphorn 2012; Chen & Stuphorn 2015; Kawaguchi et al., 2015; Seo and Lee, 2009; Lee et al., 2012), identified new types of signals related to feedback and reward processing, and revealed the laminar distribution of different kinds of signals. These observations were complemented by the first description of laminar current density associated with error and reward processing. We will discuss the nature of the various modulation patterns, new insights offered for the cortical microcircuitry of performance monitoring, and the importance for identifying cortical source(s) of the error-related and feedback-related event-related potentials.

### Signaling error, loss and gain through facilitation and suppression

Being the first sample of single-unit activity across all layers of SEF, we now relate our findings to previous reports. Overall, most neurons increased discharge rates following errors, negative feedback or lack of reward and decreased discharge rates following positive feedback and fluid reward, which agrees with previous studies in humans that report primarily signals related to negative outcomes in supplementary motor area (Mars et al. 2005). Other studies have identified more neurons signaling reward gain (Stuphorn, 2015; Lee et al., 2012), which may be due to task differences or differential sampling of neurons across layers. The magnitudes of spike rate modulation, LFP polarization and synaptic current sinks immediately after errors were all higher, the greater the cost of the error. Likewise, following feedback and during reinforcement LFP polarization and current sinks scaled with reward magnitude; however, only the magnitude of spike rate suppression in this interval scaled with gain, a pattern opposite typical dopamine neurons (Schultz 2017). Distinct neurons in our sample preferentially represented reward gain versus loss. Replicating previous reports (cited above), some neurons had elevated or reduced discharge rates more following the negative outcome cue, and others discharged more following the positive outcome cue. We also encountered neurons suppressed following reward gain, many of which were also excited for reward loss, similar to modulation in habenula (Mosamoto and Hikosaka, 2009). These unusual neurons were found almost exclusively in L2/3.

### Toward a microcircuit for performance monitoring

That elementary operations, such as predictive coding, are performed by cortical microcircuits is undisputed (Bastos et al. 2012), although agranular cortical areas have unique circuitry (Shipp 2005; Weiler et al. 2008; García-Cabezas and Barbas, 2014; Godlove et al. 2014; Beul and Hilgetag, 2015) that may be suited for performance monitoring operations (Cohen 2014). Relative to granular areas, agranular areas are distinguished by an absence of interlaminar inhibitory connections (Katzel et al. 2011). Accordingly, relative to primary visual cortex, L2/3 and L5/6 are more independent in SEF (Ninomiya et al. 2015). Consistent with this, most synaptic currents we observed associated with errors, cues and reward were sustained within layers. Inhibitory processes in L2/3 of SEF are more complex than those in L5/6 with calretinin neurons concentrated nearly exclusively in upper L2, calbindin neurons densest in L2 relative to L5/6, and parvalbumin neurons more uniformly distributed from L2 to L6 (Godlove et al. 2014). This results in another salient feature of agranular cortex ‐‐ strong, balanced excitatory and inhibitory recurrence in L2/3 whereby signals from diverse cortical and thalamic inputs are selected and integrated to drive L5/6 outputs. Consistent with this, the strongest synaptic currents we observed were centered in L2/3. Reciprocal excitation from deeper to superficial layers is weaker but hypothesized to be crucial for core operations underlying performance monitoring (Cohen 2014). The general concordance of our findings with descriptions of agranular circuitry indicates that cortical microcircuit models of performance monitoring can be formulated and tested rigorously in light of specific details understood about saccade circuitry.

As observed previously (Godlove et al. 2014), spikes wider than 250 μs increased in width with depth, consistent with arising from pyramidal cells. Spikes narrower than 250 μs did not vary in width with depth, consistent with arising from parvalbumin interneurons. Notably, error and gain neurons had wider spikes suggesting that they arise from projection neurons. Meanwhile, loss neurons had both wider and narrower spikes suggesting that they arise from both pyramidal and interneurons. Thus, only loss neurons can be postulated as inhibitory interneurons.

Knowledge of the extrinsic connectivity of SEF (Huerta & Kaas, 1990; Shook et al. 1991; Schall et al. 1993) offers functional insights into possible sources and influences of the laminar-specific signals we observed (Fig. 5). SEF receives signals about saccade production directly from FEF and indirectly from the superior colliculus through the thalamus. FEF terminals are concentrated in L2-5 of SEF, originating mainly from L2/3 pyramidal cells. SEF can receive signals about task context, plans, and rules from dorsolateral prefrontal cortex areas 12 and 46. Such representations provide a substrate for comparison of actual with planned behavior that can be detected through synaptic integration in apical dendrites in L2/3 shown by stronger current sinks on errors that manifest in error-related spiking in L2/3 and L5/6. SEF can receive signals about reinforcement and arousal from basal ganglia as well as from dopaminergic (Gaspar et al. 1992; Williams and Goldman-Rakic 1988) and noradrenergic (Aston-Jones and Cohen 2005) inputs, which can enable modulation of error-related modulation by magnitude of loss, responses to reinforcement cues and to reward delivery that signal gain and loss.

On correct and error trials a pronounced current sink was observed in L2/3 coinciding with but unrelated to mere saccade production (Godlove et al., 2014), which was significantly stronger on error trials, primarily in L3. An error can be derived by comparing a representation of a planned movement with the actual movement (Coles et al., 2001; Ridderinkhof et al., 2004). SEF receives saccade planning information via inputs from mediodorsal thalamus, which can convey efferent copy signals from SC (Sommer and Wurtz, 2008). Connectivity of DLPFC with L2/3 of SEF can support the representations of context, goal and plan (Gehring & Knight, 2000; Kerns et aI. 2004) from which an explicit error signal can be derived through coincidence detection by L2/3 neurons conveyed to L5/6 (Cohen 2014). Following early error-related spiking in deep L3 and L5, later error-related spiking occurred in L2/3 and lower L6 coincident with current sinks in L3 and L5. This sequence of spiking activation and current density corresponds roughly to that predicted by the canonical cortical microcircuit (Cohen 2014).

The laminar and temporal pattern of synaptic current and spiking modulation varied between rewarded and unrewarded trials in the cue and reward period. Coinciding with the feedback cue in correct trials was a current sink in L5/6 followed by a brief sink in L2/3. Coinciding with reward delivery was a weaker sink in L5. The rare gain neurons were recruited mainly in L3 and L5/6 after the first sink ended, consistent with dendritic integration preceding spike production. Facilitated gain neurons continued to be recruited such that this signal persisted after the reward was consumed. However, the sample of suppressed loss neurons, which were mainly in L2/3, were notably less active immediately before and after the reward was delivered. Intralaminar inhibition is a likely source of this suppression. The gain neurons and many suppressed loss neurons had wide spikes consistent with projection neurons. Thus, many distant cortical and subcortical SEF efferent targets can be informed of task success while output to cortical areas involved in exerting adjustments in behavior are suppressed.

In unrewarded error trials, the first sink was later and more superficial in L3 or L5, and the other two sinks were replaced by a weak, brief sink in L6. When reward would have been delivered, no sinks were observed. Meanwhile, the many loss neurons were recruited predominantly in L2 and L6 plus L3 when the initial sink ended. Some loss neurons had narrow spikes consistent with being inhibitory parvalbumin neurons. Facilitation of such loss neurons would impose inhibition on the local circuit within the layers. Suppression would release inhibition, facilitating post-synaptic neurons. We observed differential current density across high and low loss trials that was sustained within layers. This is consistent with inhibition acting within rather than across layers. After the early surge of response suppression, neurons in all layers were both facilitated and suppressed, providing the opportunity for signals about positive reinforcement to be delivered to cortical and subcortical efferent targets.

Thus, our new findings offer insights into how SEF can monitor performance. Our findings also offer insights into possible contributions of SEF to executive control. Spiking and LFP modulation has been found in ACC signaling errors and reward during this task (Ito et al. 2003; Emeric et al. 2007, 2010). Of note, error-related modulation in ACC followed that in SEF, suggesting that neurons in SEF signaling error and loss convey this signal to ACC. Via this pathway, processes in the locus coeruleus can be influenced (Kalwani et al. 2014). SEF can influence saccade production directly through efferents to FEF (with terminals predominantly in L1-4 originating from pyramidal cells in L2/3 and deep L5), to SC (with terminals in the intermediate and deep layers), and to omnipause neurons in the nucleus raphe interpositus. Drawing two pathways from SEF to FEF in Fig. 5 emphasizes the fact that projections from different layers are likely to convey different signals. Previous work has demonstrated that subthreshold electrical stimulation of SEF improves saccade countermanding performance by delaying RT (Stuphorn and Schall 2006), and adaptive RT slowing is accomplished by postponing the beginning of accumulation of presaccadic activity in FEF and SC (Pouget et al. 2011). These anatomical and functional relationships point to the hypothesis that RT slowing is enabled by projections to the saccade circuitry by loss neurons in L2/3 and error neurons in L5. Strategic adaptation of RT can also be enabled by projections of SEF to striatum (Parthasarathy et al. 1992) where neurons convey reinforcement signals (Hikosaka et al. 1989). In L6 we observed a high frequency of error and loss neurons, which can influence processing of corollary discharge, which is disrupted in SZ patients (Thakkar et al. 2015).

Although omitting known connections of SEF, this survey highlights the tractability of elucidating a microcircuit level model of performance monitoring. Such a model requires filling a number of specific gaps in our knowledge. Fortunately, methods are available to obtain major elements of the required information, which can guide the next generation of cortical microcircuit model for medial frontal cortex. Such models can be firmly grounded on interactive race models of countermanding performance (Boucher et al. 2007; Lo et al. 2009) and related to network level models (Wiecki and Frank 2013) to rationalize therapies for psychopathologies.

### Seeking the source of error and feedback event-related potentials

Finding greater current density in SEF on error as compared to correct trials confirms that SEF contributes to the ERN, in contrast to the suggestions of some authors (Cole et al., 2009; but see Schall & Emeric 2010). Located on the dorsomedial convexity, SEF is ideally positioned to contribute to voltage polarizations recorded over medial frontal cortex, and our finding of stronger synaptic currents in L2/3 relative to L5/6 is consistent with observations in human ACC (Wang et al. 2005). Neural discharges and synaptic current flow coincided with intervals when ERN and FRN occur, and various characteristics of the SEF signals paralleled previous observations for the ERN/FRN, e.g., sensitivity to cost of error (Gehring et al. 2012). However, the differences in latency, layer and polarization observed with error detection contrasted with gain/loss feedback indicate that the ERN and FRN have the different neural generators within a single cortical area.

### Conclusion

Deep insights into the microcircuitry of primary visual cortex began with studies elucidating the properties of neurons in different layers (Gilbert 1977; Leventhal and Hirsch, 1978) and descriptions of laminar patterns of current flow (Mitzdorf and Singer, 1979). This study provides the first equivalent information for the SEF. Being an agranular area, comparisons and contrasts with primary sensory areas provide insights into the degree of uniformity of cortical areas. As a likely source contributing to the ERN/FRN, details about laminar processing in SEF offer unprecedented insights into the microcircuitry of performance monitoring.

## Methods

This paper is based on an analysis of data collected previously (Godlove et al, 2014), so we will only summarize essential information here before elaborating the new analyses. All procedures were approved by the Vanderbilt Institutional Animal Care and Use Committee in accordance with the United States Department of Agriculture and Public Health Service Policy on Humane Care and Use of Laboratory Animals.

Data were collected in two head fixed macaque monkeys (monkey Eu Macaca radiata and monkey X Macaca radiata) performing a memory-guided saccade task and a countermanding task (Logan and Cowan, 1984; Hanes and Schall 1995). The countermanding task was performed with two target locations (Fig. 1). For this study one target location was associated with larger magnitudes of fluid reward than the other location. The lower magnitude reward ranged from 0-50% of the higher magnitude reward and was adjusted to encourage the monkey to continue responding to both targets. The location of the high reward target changed across blocks of trials. The number of trials in each block was determined by the number of correct trials performed. Block length was adjusted to maintain performance at both targets. In most sessions block length was set at 10 to 30 correct trials. As in previous implementations of asymmetrically rewarded tasks (Kawagoe et al., 1998), errors led to repetitions of target location, ensuring that monkeys did not neglect low-reward targets in favor of high-reward targets.

Using methods detailed before (Godlove et al. 2014), spiking activity and local field potentials were recorded in SEF from five sites with a 24-channel linear electrode array with 150 μm spacing (Plexon U-probe). Three penetrations were oriented perpendicular to the cortex at sites from which saccades were evoked with low currents. Neural recordings were assigned to specific cortical layers using the approach described previously (Godlove et al. 2014; see Supplementary Methods and Supplementary Fig. 2).

Patterns of spiking activity were classified according to criteria similar to those described previously (Stuphorn et al. 2000; Stuphorn et al. 2010). Baseline discharge rate was defined as the mean spike density with the corresponding standard deviation of 300 ms prior to target onset. Trials were aligned on saccade initiation for analysis of error-related activity and on auditory feedback tone for analysis of reinforcement (and feedback)-related activity. Error‐ and reinforcement-related spiking activity were defined by the differential activity between postsaccadic discharge rate on error and correct trials, and post-tone discharge rate on rewarded and unrewarded trials, respectively (see Supplementary Methods). Field potentials were low-pass filtered (<30Hz) (Emeric et al., 2010) and CSDs were calculated (see Supplementary Methods). Grand average error‐ and reinforcement-related polarization in field potential and current source density were obtained by normalizing these signals across sessions and a pixel-by-pixel comparison relative to a 200ms baseline (Wilcoxon test, see Supplementary Methods). The differential LFPs and CSDs for error‐ and reinforcement-related analyses were calculated based on the difference between those on correct and error trials (aligned on saccade onset), and difference between rewarded and unrewarded trials (aligned on feedback tone) respectively.

